# Dynamical Regimes in Rejuvenation

**DOI:** 10.64898/2026.08.27.747604

**Authors:** Matteo Ciarchi, Steffen Rulands

## Abstract

Biological aging is accompanied by systematic changes in epigenetic modifications and chromatin organization. The reversal of the effects of aging, rejuvenation, is experimentally achieved by the transient induction of factors that modify these marks in cells and organisms. Here, we show that key features of rejuvenation experiments emerge from the biophysical interplay between dynamic epigenetic marks and the three-dimensional conformation of chromatin. Using a minimal field theory and molecular dynamics simulations, we show that the system responds in three distinct temporal regimes. The intermediary regime fulfills necessary conditions for successful rejuvenation. In this regime, the system spends time near a separatrix, allowing for high epigenetic plasticity, while memory retained in the chromatin conformation enables restoration of the original epigenetic correlations. Analysis of sequencing data further supports the predicted coupling between chromatin compaction and epigenetic correlations. Our results provide a physical explanation for how rejuvenation may remodel age-associated epigenetic states without irreversibly erasing cellular identity. We identify a general mechanism by which memory stored in a slow structural variable permits reversible remodeling of a faster internal state.

---

Cells must stably maintain their identity while responding rapidly to specific signals [1–6]. At the molecular scale, this duality is reflected in both dynamic epigenetic modifications, including DNA methylation and histone modifications, and the three-dimensional folding of chromatin in the cell nucleus [7]. During aging, epigenetic states and chromatin organization undergo progressive remodeling [8–11]. Machine-learning models use the tight correlation between DNA methylation and age to estimate biological age and mortality risk with high accuracy [12, 13].

Consequently, recent experimental approaches to rejuvenation target these epigenetic marks. These rejuvenation strategies are largely based on transient cellular reprogramming experiments in which cells or animals are exposed to the transcription factors Oct4, Sox2, and Klf4 (OSK), with some protocols additionally including c-Myc (OSKM) [14–17]. A central observation is that outcomes depend sensitively on the duration and scheduling of OSK(M) induction [15, 17, 18] [Fig.1(a)]: exposure must be transient or cyclic to remodel age-associated features while limiting loss of cellular identity [19, 20]. Very short exposures are ineffective, whereas prolonged expression can lead to an irreversible loss of cell identity. These observations suggest that successful rejuvenation requires a transient state in which age-associated epigenetic features become remodelable while sufficient information about the somatic state remains collectively stabilized.

Here, we show that the biophysical interplay between dynamic epigenetic marks and the conformation of the DNA and chromatin in space is sufficient to understand key features of rejuvenation experiments: the response in three distinct temporal regimes and the simultaneous emergence of plasticity and memory in the intermediate regime. Specifically, we define a minimal field-theoretical model coupling chromatin conformation in three-dimensional space to the dynamics of epigenetic marks, accounting for both the local chromatin dependence of mark deposition and the reciprocal effect of these marks on chromatin compaction [7, 21–23]. We derive the phase behaviour of the model and study its behaviour in response to temporal driving. Our results are validated using molecular dynamics simulations. This identifies a general physical mechanism by which the epigenome can become transiently plastic without erasing its initial state.

## Chemical-conformational field theory

To test whether biophysical properties of the epigenome and chromatin can predict the temporal response regimes observed in rejuvenation experiments, we define a minimal field-theoretic description of the interaction between chromatin structure and epigenetic marks [Fig. 1(b)]. Epigenetic marks can be categorized into active marks, such as H3K4me3, which are positively associated with gene expression, and repressive marks, like H3K27me3, that are involved in gene silencing. Active and repressive marks can compete for local chromatin occupancy and influence chromatin conformation [24–26]. A minimal model therefore comprises the competition between active and repressive modifications along the one-dimensional genome and their interaction with the three-dimensional conformation of chromatin in space [27–31]. We first characterize the equilibrium states of the coupled system and subsequently introduce relaxational dynamics to describe its response to time-dependent perturbations. The equilibrium configurations minimize the total free energy *ℱ* comprising contributions from chromatin conformation, the epigenetic field, and their interaction.

**Figure 1.**
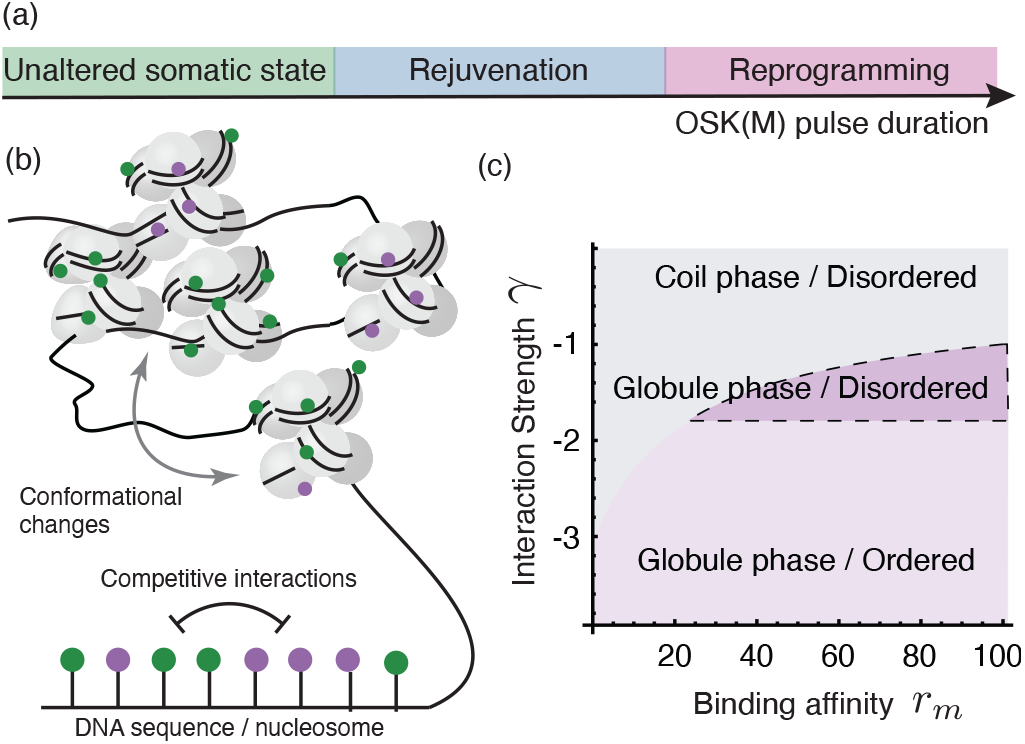
(a) Three temporal regimes in rejuvenation experiments. (b) Sketch of the model comprising the interplay between interacting epigenetic marks and the conformation of chromatin. (c) Phase diagram of the model as a function of the interaction strength *γ* and the enzyme binding affinity *r*_*m*_, fixing *u*_2_ = 3.0, *u*_3_ = 1.0, *λ* = 0.3, *µ* = 0.5. Shaded areas signify different phase behaviours.

## Phase behavior

Before studying the effect of time-dependent rejuvenation protocols, we first consider the equilibrium and response behavior of the model. We ask whether the system can support stable long-range order, representing memory, while remaining responsive to perturbations that rewrite epigenetic marks. This feedback involves the mean epigenetic field and the two-point correlation functions of *m*(*x*) and 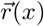. With these quantities, memory is reflected in the existence of multiple stable fixed points of the ensemble average *M* = ⟨*m*(*x*)⟩. Here, *M* is a coarse-grained order parameter which specifies the genomic average of the modeled epigenetic imbalance, but not its spatial distribution or the complete molecular epigenetic state of a cell. To be stable, these fixed points must also exhibit long-range correlations in the epigenetic field, ⟨*m*(*x*)*m*(*x*′)⟩ ∝ |*x* − *x*′|^−*β*^, with *β <* 2.

To study the phase behavior of the system, we use the two-particle-irreducible (2PI) effective-action formalism [32, 33], which yields self-consistent equations for the connected propagators *G*_*m*_ and *G*_*r*_ and resums infinite classes of perturbative contributions ([34]). Crucially, because *G*_*m*_ and *G*_*r*_ appear as independent variational objects, the formalism naturally encodes the mutual back-reaction between chromatin conformation and epigenetic correlations without imposing any a priori decoupling between the two modalities.

Using this approach, we derive stationarity conditions for *G*_*r*_, *G*_*m*_, and *M* in periodic boundary conditions (chain polymer). The full stationarity condition for the correlator describing chromatin conformation, *G*_*r*_, is shown in [34]. To first order in *γ, u*_2_, and *u*_3_ and in the limit *q* → 0, substitution of the Fourier-space ansatz 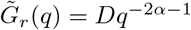 gives

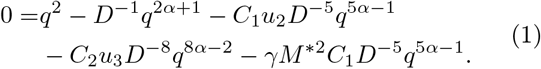

Here, the bare second and third virial coefficients, *u*_2_ and *u*_3_, measure epigenetic-state-independent interactions between pairs of chromatin segments. The coupling *γ <* 0 measures the energetic stabilization of contacts between chromatin segments carrying the same epigenetic state and therefore quantifies the strength of contact-mediated reader-writer feedback.

Equation (1) alone reproduces the three known phases of an interacting polymer in the mean-field approximation: the self-avoiding coil (*u*_2_ *>* −*γM*^∗2^, *α* = 3/5), the ideal Gaussian chain (*u*_2_ = −*γM*^∗2^, *α* = 1/2), and the collapsed globule (*u*_2_ *<* −*γM*^∗2^, *α* = 1/3). Importantly, the effective two-body interaction is renormalized by the epigenetic mean field *M*^∗^, so that the boundaries between polymer phases are shifted by the average of the epigenetic field. Similarly, we obtain stationarity conditions for the ensemble average of the epigenetic field, *M*,

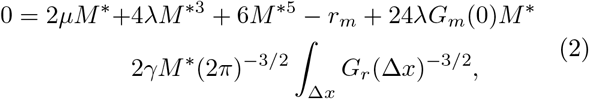

and its correlator *G*_*m*_,

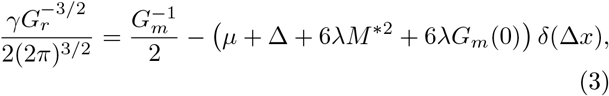

where Δ*x* = *x* − *x*′. The parameter *µ* sets the intrinsic free-energy cost of establishing a local excess of one class of epigenetic marks over the other. *λ >* 0 and the sextic term account for the finite occupancy of chromatin and prevent the local mark imbalance |*m*| from growing without bound. Finally, *r*_*m*_ is the uniform binding affinity of methylating enzymes. Together, these conditions reveal the reciprocal dependence between chromatin conformation and the epigenetic field: the large-distance behavior of *G*_*m*_ is governed by 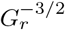, which is the contact probability in the Gaussian and collapsed phases. In these cases, this dependence induces long-range epigenetic correlations scaling as |*x* − *x*′| ^−3*/*2^ and |*x* − *x*′|^−1^, respectively, allowing for phase transitions in the epigenetic field driven by chromatin organization [35]. In the self-avoiding phase, fluctuation corrections suppress the contact probability, which then scales approximately as |*x* −*x*′|^−2.1^ [36], sufficiently rapidly that epigenetic interactions become effectively short-ranged, precluding long-range order.

To characterize the phase behavior of the coupled chromatin–epigenetic system, we solve the self-consistent equations (1) and (2) numerically. The resulting phase diagram contains three regimes distinguished by polymer conformation, mean value of epigenetic field, and its correlations along the genomic contour (Fig. 1c). Consistent with Ref. [27], we find three phases: one, for weakly interacting epigenetic marks, with coiled polymer conformation, disordered *m*(*x*) (single stable equilibrium for *M*) and short-ranged correlations, and a second phase with compact (globule) polymer conformation where two stable nonzero values coexist with long-ranged correlations *G*_*m*_, producing robust ordered epigenetic attractors. Our analysis also revealed a third phase, where the polymer adopts a compact globular conformation and long-range correlations in *m*(*x*) but with a single stable equilibrium with *M* ≠ 0.

## Comparison to experimental data

Equations (1) and (3) predict that a compact chromatin domain should exhibit more slowly decaying epigenetic correlations than an open domain. We therefore expect the connected correlation function of a repressive mark to decay more slowly in compact B compartments than in open A compartments, with a smaller compartment-dependent difference in cells with weak compartmentalization. To test this, we calculated the connected correlation function of H3K9me3 from ChIP-seq measurements in the A and B compartments of mouse embryonic fibroblasts (MEFs) and 2i mouse embryonic stem cells (2i ESCs) [37–39]. Because ChIP-seq is nonstoichiometric, the comparison is qualitative and concerns correlation range rather than absolute amplitude. MEFs have well-defined compartments, with A predominantly open and active and B more compact and repressive [40, 41]. The H3K9me3 correlation function decays more slowly in B than in A [Fig. 2, left], consistent with the predicted conformation dependence of *G*_*m*_. In 2i ESCs, which have less condensed chromatin and weaker compartmentalization [39, 42, 43], the A- and B-compartment correlations are more similar and the distinct long-distance regime of the MEF B compartment is absent [Fig. 2, right]. These observations do not test rejuvenation or establish causality. They are consistent with the predicted association between compact chromatin and a more slowly decaying *G*_*m*_.

**Figure 2.**
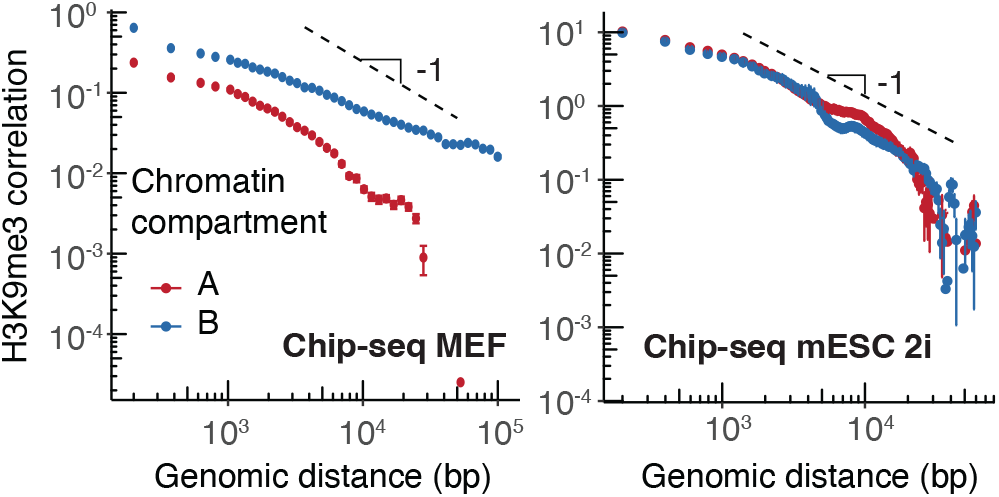
(a) Connected correlation functions of H3K9me3 in MEFs and mESCs under 2i conditions, grouped by chromatin compartment (A, B). Points represent averages over 50 bins along the *x* axis, error bars indicate the standard error. The dashed line represents the scaling predicted by the theory for a fractal globule at large distances. Because ChIP-seq is nonstoichiometric, the comparison between theory and experiment is only qualitative.

## Epigenetic dynamics in rejuvenation

To study the dynamics, we derive equations for the coevolution of the mean epigenetic field *M* and the mean extension 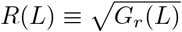 of a genomic region of contour length *L* by taking functional derivatives of the effective action at zero binding affinity *r*_*m*_ = 0 [34]. This defines two characteristic time scales, *T*_*R*_ and *T*_*M*_, describing the typical global relaxation time of the polymer and the epigenetic field, respectively. These are determined by the inverse mobilities and the local curvatures of the effective-action landscape (see [34]).

The resulting phase portraits provide a reduced representation of the field dynamics in terms of these two collective coordinates. Each point in the (*R, M*) plane therefore represents many microscopic configurations of chromatin and epigenetic marks, while the separatrices describe basin boundaries within this reduced two-variable model. In the coiled phase, the dynamics is governed by a single fixed point with *M* = 0 and self-avoiding polymer statistics; this state has open chromatin and lacks collective epigenetic memory. In the globule phase [Fig. 3(a)], two additional stable fixed points appear, corresponding to the predominance of either epigenetic mark. Their local stability prevents uncorrelated fluctuations from erasing information stored in the epigenetic field, thereby providing long-term *epigenetic memory*. Therefore, changes in chromatin conformation permit both epigenetic memory [23] and plasticity [6].

**Figure 3.**
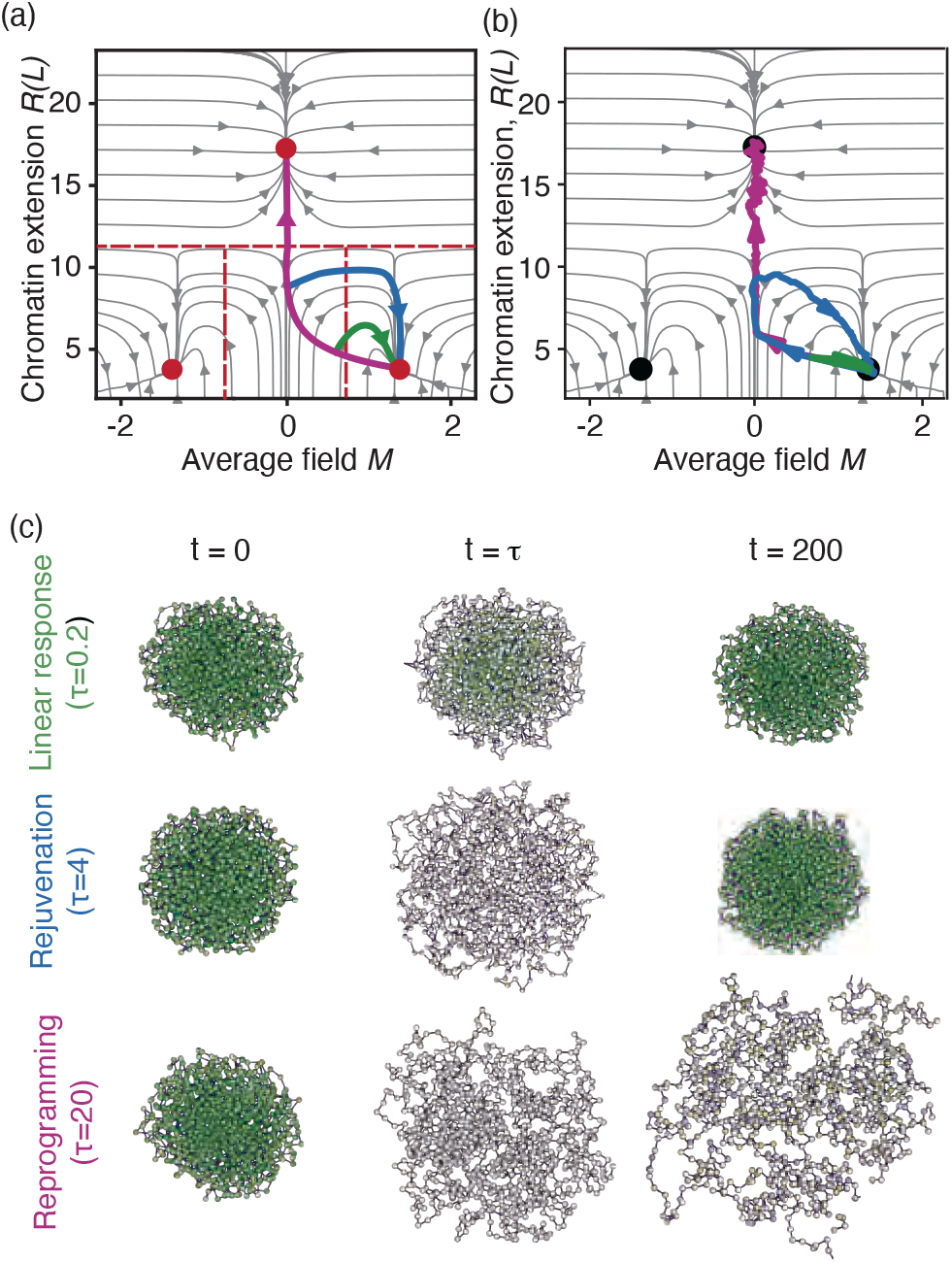
(a) Flow field for the polymer extension *R*(*L*) and average epigenetic field *M* for *γ* = −1.5. Fixed points are indicated by red dots. Sample trajectories under three pulse durations for *T*_*M*_ *< T*_*R*_ are sketched: green short, blue intermediate, and purple long. (b) The trajectories from three Langevin dynamics simulations are superimposed on the phase portrait in (a) (*τ* = 0.2 green, *τ* = 4.0 blue, and *τ* = 20 purple) (See [34] for details) (c) Snapshot configurations of the same simulations shown in (b). Green and purple colors indicate the predominance of one of the two epigenetic marks.

We now investigate how these biophysical properties affect the response of chromatin conformation and epigenetic marks under temporal driving in rejuvenation experiments. We assume that, in these experiments, OSK(M) acts primarily on the epigenetic variable rather than directly on chromatin conformation. We consider a genomic domain in which the average value of *m*(*x*) rests at one of the fixed points in the globule phase. OSK(M) induction is then represented as a minimal erasure of somatic epigenetic order. We describe its effect by an external field *h*(*x, t*) acting on *m*(*x*) in such a way that it reduces the absolute value of *M* by modifying the free energy as ℱ + ∫_*x*_ *h*(*x, t*)|*m*(*x*)|. We consider constant OSK(M) induction over a period *τ*, *h*(*x, t*) = *h*_0_ [Θ(*t*) − Θ(*t*−*τ*)].

The phase portraits in Fig. 3(a) provide an interpretive framework for transient reprogramming experiments. The effect of the reprogramming protocol is determined by the relative strength of the effective stimulus *h M*^∗^ compared with other energy scales. The trajectory in phase space during transient reprogramming depends both on the absolute strength of the stimulus, *h*, and its duration, *τ*. In particular, the dynamical regimes are controlled by the epigenetic inverse mobility and chromatin relaxation times, Γ_*m*_ and *T*_*R*_, the positions of the separatrices in the (*R, M*) plane defined by Eqs. (1) and (2), and the strength of the external stimulus.

We consider escape from a stable fixed point *M*^∗^. We focus the analysis on the regime in which epigenetic remodeling occurs faster than the large-scale conformational response, corresponding in the strong-driving limit to *hT*_*R*_/Γ_*m*_ *> M*^∗^ [Fig. 3(a)]. Such a dynamical ordering is consistent with time-resolved reprogramming experiments, in which histone-mark remodeling can precede A/B compartment switching and substantial biochemical remodeling occurs while target regions remain embedded in a highly interacting chromatin environment [44, 45]. In the fast-epigenetic regime, switching on the reprogramming field first drives a rapid relaxation of *M* at nearly fixed chromatin conformation. Thereafter, the trajectory follows the driven nullcline ∂*M*/∂*t* |_*R*_ = 0 after a rapid transient.

Relaxation after the pulse then falls into three regimes determined by the magnitude of the epigenetic perturbation *hτ*. For short-term OSK(M) exposure, 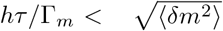, the strength of the stimulus is comparable to the strength of fluctuations in *m*(*x*). Both mean order and long-distance correlations remain close to their initial values. The subsequent dynamics is described by linear-response theory, and relaxation is governed by the stable attractor nearest the initial compact-chromatin state. Biologically, the epigenome converges to its original configuration.

For intermediate exposure times, *hτ*/Γ_*m*_ ≈ *M*^∗^, the system is driven along the nullcline defined by ∂*M*/∂*t*|_*R*_ = 0. The trajectory approaches the separatrix *M* = 0 while remaining on the compact side of the globule–coil boundary. Mean order is suppressed, producing plasticity, whereas the algebraic component of *G*_*m*_ preserves correlated structure. We identify this memory-preserving neighborhood of the separatrix as the candidate rejuvenation regime. In the reduced (*R, M*) phase portrait, this separatrix is the boundary between the basins of the two ordered epigenetic states and therefore represents a plastic regime with high susceptibility in which small perturbations can redirect the dynamics. In addition, since the polymer itself remains in a globule state, in this regime memory about the configuration of epigenetic marks along the genome is retained geometrically and can be recovered in the epigenetic field *m*(*x*) after relaxation.

Finally, for long-term OSK(M) exposure, *τ* ≳ *M*^∗^Γ_*m*_/*h* + *T*_*R*_, the system undergoes a globule-to-coil transition. Once the trajectory crosses the globule–coil boundary, the contact-mediated interaction becomes effectively short-range. The loss of the algebraic component of *G*_*m*_ removes the correlational memory that distinguishes the intermediate regime. After the end of exposure, the system relaxes to an open, epigenetically disordered fixed point.

Together, these results resemble key characteristics of rejuvenation experiments. The response to OSK(M) follows three temporal regimes set by the stimulus and by the separation between epigenetic and polymer relaxation times. The intermediary regime fulfills necessary criteria for rejuvenation: plasticity and memory of the original somatic state.

## Molecular dynamics simulations

To test these predictions numerically, we performed Langevin molecular dynamics simulations of a bead-spring polymer of *N* monomers, each carrying a continuous scalar *m*_*i*_ that represents the local balance between two epigenetic states. Reprogramming is modeled by applying an external field *h* coupled to |*m*_*i*_| for a duration *τ*, after which the field is removed. See [34] for details. The simulations reproduce the three predicted dynamical regimes [Fig. 4]. In the intermediate regime, the epigenetic susceptibility *χ* is enhanced (Fig. 4(a)) while the locus-specific chromatin conformation retains substantial overlap with its initial state [Fig. 4(b)]. Remarkably, the epigenetic overlap vanishes near the separatrix but reappears during relaxation [Fig. 4(c)], consistent with the hypothesis that information retained in chromatin conformation can restore the original epigenetic correlations up to global sign inversion.

**Figure 4.**
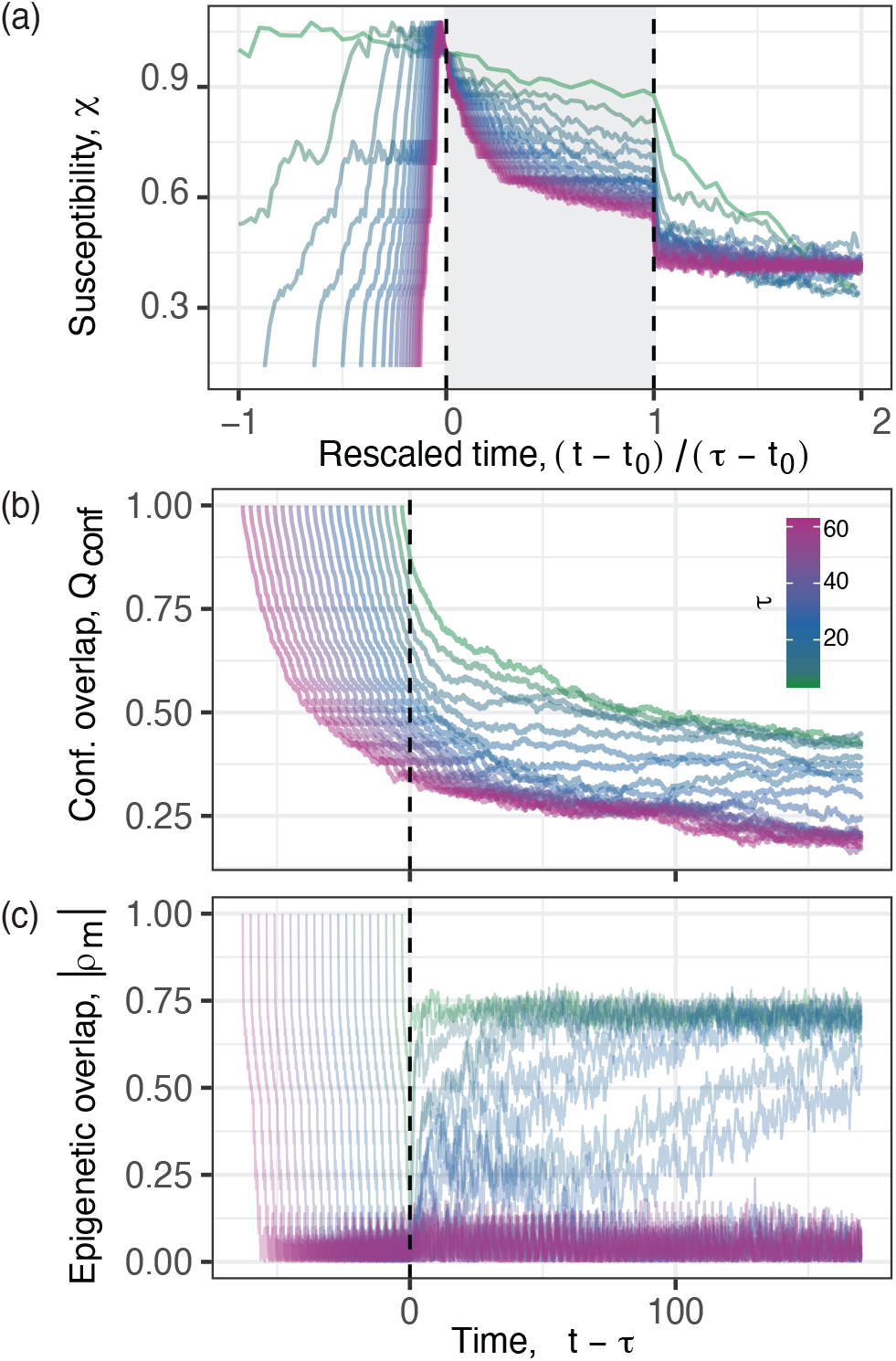
Simulation results for a polymer of *N* = 512 beads at different protocol durations (colorbar). (a) Response to a short-term external field (susceptibility) over time. (b) Conformational overlap of the polymer compared to its conformation at *t* = 0. *t*_0_ is the time the trajectory approaches the separatrix (|*M*|*<* 0.05). The shaded area represents time spent near the separatrix. (c) Overlap of fluctuations in *m*(*x*) compared to *t* = 0. Dashed lines in (b) and (c) represent pulse withdrawal. Mathematical definitions and simulation parameters are in [34].

## Discussion

We have shown that a minimal model comprising interactions between chromatin conformation and epigenetic dynamics captures key characteristics of current rejuvenation experiments: As in experiments, the system responds in three distinct temporal regimes. The intermediary regime fulfills necessary conditions for successful rejuvenation. In this regime, the system spends time on a separatrix, allowing for epigenetic plasticity, while conformational memory enables restoration of the original epigenetic correlations. That said, the construction of a biophysical theory does not comprise the full complexity of epigenetic regulations: it cannot explain which features of the epigenome lead to functional rejuvenation or how the symmetry between the two somatic fixed points is broken biologically. Our results predict that perturbations of chromatin compaction or reader–writer feedback will shift the induction times separating reversible remodeling, plasticity, and loss of identity. Time-resolved measurements of chromatin contacts and multiple epigenetic marks during OSK(M) induction could directly test this prediction.

We thank V. M. Schimmenti for help in developing the code used for the molecular dynamics simulations, and Peng Rao for useful discussions on the field theory. S.R. and M.C. received funding from the European Research Council (ERC, grant agreement no. 950349). S.R. is a member of the Center for NanoScience (CeNS).

## Declaration of AI usage

chatGPT 5.6 was used for assistance with coding and text editing. All outputs were reviewed and verified by the authors.

## Notes

### Competing Interest Statement

The authors have declared no competing interest.

### Summary of Updates

Author list: Corrected order of author list, such that Matteo Ciarchi is first author and Steffen Rulands is last author in the online form.

