## Supplementary Material for "Dynamical Regimes in Rejuvenation"

#### S1. MODEL DEFINITION

In this section, we introduce the field theory of the coupled epigenetic–chromatin system. Briefly, at a given genomic position $i$ we define two binary variables, $s_i^1$ (repressive, e.g. H3K9me3) and $s_i^2$ (active, e.g. H3K4me3/H3K27ac). Both kinds of marks tend to be mutually exclusive and undergo competitive dynamics [24–26]. The epigenetic state at a given position is then summarized in a variable $m_i = s_i^1 - s_i^2$, such that positive values of $m_i$ correspond to a local enrichment of repressive marks and negative values to a local enrichment of active marks. Following the Ginzburg-Landau argument, we expand the free energy part of the epigenetic scalar field $m(x)$ in lowest order and in the continuum limit. The free energy contribution of the epigenetic $m(x)$ field then takes the form

$$\mathcal{F}_m = \int_x \left[ (\partial_x m(x))^2 + \mu m(x)^2 + \lambda m(x)^4 + m(x)^6 - r_m m(x) \right]. \tag{S1}$$

The gradient term, as the lowest-order derivative, reflects that interactions between epigenetic marks can extend spatially along the genome [48, 49]. We discard this term in the current analysis because it does not affect the long-distance scaling properties of $m(x)$. The parameters $\mu$ and $\lambda$, and the term $m(x)^6$ give the strengths of demethylation processes and site exclusion, respectively. The external field $r_m$ represents a binding affinity for the epigenetic marks that biases the average of $m(x)$ toward high values when positive.

We describe chromatin conformation using the Edwards model, the continuum limit of a polymer chain with excluded-volume interactions. This model has been used to describe chromatin organization on the scale of kilobases [40, 46]. The chromatin part of the free energy is described by the continuum model of a self-avoiding walk. Its contribution to the free energy reads

$$\begin{aligned}\mathcal{F}_\mathrm{p} = &\int_x (\nabla \vec{r}(x))^2 + u_2 \int x, x' \delta(\vec{r}(x) - \vec{r}(x')) \\ &+ u_3 \int_{x,x',x''} \delta(\vec{r}(x) - \vec{r}(x'))\delta(\vec{r}(x') - \vec{r}(x'')),\end{aligned} \tag{S2}$$

where we denote the position of genomic locus $x$ in three-dimensional nuclear space by $\vec{r}(x)$. The first term corresponds to the entropic contribution to chain elasticity, and the second and third terms represent, respectively, two- and three-body interactions [32], with $u_2$ and $u_3$ positive constants describing the second and third virial coefficients. The final component of the model is the interaction between epigenetic modifications and chromatin conformation. This interaction favors conformations in which chromatin sites carrying the same epigenetic state are close in three-dimensional space. Therefore, we introduce a free-energy term that is minimized when same-sign modifications are in spatial proximity:

$$\mathcal{F}_\mathrm{int} = \gamma \int_{x,x'} m(x)m(x')\delta(\vec{r}(x) - \vec{r}(x')). \tag{S3}$$

The parameter $\gamma < 0$ controls the strength of this feedback.

The reprogramming protocol is modeled by a modification of the free energy that disfavors the presence of any mark. This is introduced as a field $h > 0$ conjugate to $|m|$: the coupling to the absolute value of $m$ makes any $m \neq 0$ energetically unfavorable:

$$\mathcal{F}_\mathrm{repr.} = \int_x h|m(x)|. \tag{S4}$$

A constant OSK(M) induction over a period $\tau$ starting from $t = 0$ is given by $h(x,t) = h_0 \left[\Theta(t) - \Theta(t - \tau)\right]$. We measure free energies in units of $k_\mathrm{B}T$. The continuum theory is supposed to describe the physics on long distances in sequence space. In real terms, this would correspond to distances starting from the order of a couple of nucleosomes.

### S2. 2PI EFFECTIVE ACTION FORMALISM

#### A. Diagrammatic expansion of the 2PI action at lowest order: minimization conditions

In this section, we derive the stationarity conditions for the two-particle-irreducible (2PI) effective action associated with the free energy of the coupled polymer–field system:

$$\mathcal{F} = \mathcal{F}_m + \mathcal{F}_{\rm p} + \mathcal{F}_{\rm int}. \tag{S5}$$

We introduce the 2PI effective action functional for the fields $M(x), \vec{R}(x), G_m(x,x'), G_r(x,x')$:

$$\begin{aligned}\Gamma\{M,\vec{R},G_m,G_r\} =& I(M,\vec{R}) - \frac{1}{2}\,\mathrm{Tr}\{\ln G_m\} + \frac{1}{2}\,\mathrm{Tr}\{G_{m0}^{-1}G_m\}\\ &- \frac{1}{2}\,\mathrm{Tr}\{\ln G_r\} + \frac{1}{2}\,\mathrm{Tr}\{G_{r0}^{-1}G_r\}\\ &+ \Gamma_2\{M,\vec{R},G_m,G_r\} + \mathrm{const},\end{aligned} \tag{S6}$$

where $I(M,\vec{R})$ is the tree-level free energy $\mathcal{F} = \mathcal{F}_m + \mathcal{F}_{\rm p} + \mathcal{F}_{\rm int}$, and $\Gamma_2$ collects all 2PI diagrams built from the interaction vertices of the free energy with fields $m, \vec{r}$ shifted by their means $M, \vec{R}$. The physical mean fields and propagators are then given by the stationarity conditions of Eq. (S6).

The starting point is Eq. (S6). In this expression, $I(M,\vec{R})$ is obtained by considering the tree-level part of the free energy, where the fields $m(x)$, $\vec{r}(x)$ have been substituted by the ensemble averages $\langle m(x)\rangle = M(x)$, $\langle \vec{r}(x)\rangle = \vec{R}(x)$. This gives

$$I(M,\vec{R}) = \int_x (\nabla\vec{R}(x))^2 + \int_x \left[(\nabla M(x))^2 + \mu M(x)^2 + \lambda M(x)^4 + M(x)^6 - r_m M(x)\right]. \tag{S7}$$

Since the two- and three-body interaction terms and the epigenetic field–chromatin interaction depend on nonlocal operators, it is not convenient to express them as a purely $\vec{R}$-dependent contribution to $I(M,\vec{R})$. We therefore include their shifted forms in the interacting part of the 2PI action, $\Gamma_2\{M,\vec{R},G_m,G_r\}$, in Eq. (S6). This gives, for the delta operators, contributions of the form $\delta(\vec{R}(x)-\vec{R}(x')+\vec{r}(x)-\vec{r}(x'))$. An approach avoiding the introduction of $\vec{R}(x)$ is also possible by introducing a mixed 2PI functional for $m$ and a Luttinger–Ward functional for $\vec{r}$.

The traces in Eq. (S6) are taken over the space coordinates and the vector components of $\vec{r}$. The products of the connected correlation functions $G_m(x,x') = \langle m(x)m(x')\rangle - M(x)M(x')$ and $G_r(x,x') = \frac{1}{3}\langle|\vec{r}(x)-\vec{r}(x')+\vec{R}(x)-\vec{R}(x')|^2\rangle$ are to be considered as products in operator space, e.g. $G_mG_m = \int_{x''} G_m(x,x'')G_m(x'',x')$. $G_{m0}$ and $G_{r0}$ are the free propagators of the fields in the shifted free energy. Their inverse in Fourier space (for a space-independent $M(x)=M$) is $G_{m0}^{-1}(q) = \mu + q^2 + 6\lambda M^2$ and $G_{r0}^{-1} = q^2$.

Translational invariance in three-dimensional space imposes $\vec{R}(x) = \langle\vec{r}(x)\rangle = 0$. We therefore set $\vec{R}(x) = 0$ in the following calculations for $M(x)$, $G_r$, and $G_m$. Technically, the 2PI functional is usually introduced for the correlator $\langle\vec{r}(x)\cdot\vec{r}(x')\rangle$. As the interaction vertices are given by translation-invariant operators, we will consider instead in the expression for the diagrams the form of $G_r$ introduced above. For the equation of $G_r$ itself, this amounts to discarding the IR divergent part present in $\langle\vec{r}(x)\cdot\vec{r}(x')\rangle$.

The term $\Gamma_2\{M,\vec{R},G_m,G_r\}$ is given by all 2-particle-irreducible Feynman diagrams obtained from the interaction vertices of the free energy Eq. (S5) upon shifting of the fields $m(x)$ and $\vec{r}(x)$ by $M(x)$ and $\vec{R}(x)$ respectively (the 2-particle-irreducible diagrams are those Feynman diagrams that cannot be disconnected by cutting two internal lines). In our case, these diagrams are drawn from the vertices indicated in Fig. S1.

When considering the Feynman diagrams for $\Gamma_2\{M,\vec{R},\ G_m,\ G_r\}$, solid lines represent $G_m$, the connected correlation function in the 2PI effective action. The treatment of diagrams containing polymer interaction lines is more complicated because the interaction is not polynomial in the $\vec{r}(x)$ field. This implies that there is no general substitution for the wavy line in diagrams in terms of the polymer-field correlation function $G_r$.

We consider the contributions of the Feynman diagrams to $\Gamma_2\{M,\vec{R},\ G_m,\ G_r\}$ at lowest order in the coupling constants For computational simplicity, we discard contributions coming from diagrams involving sextic interactions, as they amount to a mass shift. The $m(x)^4$ interaction produces the term

$$D_\lambda = 6\lambda\int_x G_m(x,x)G_m(x,x). \tag{S8}$$

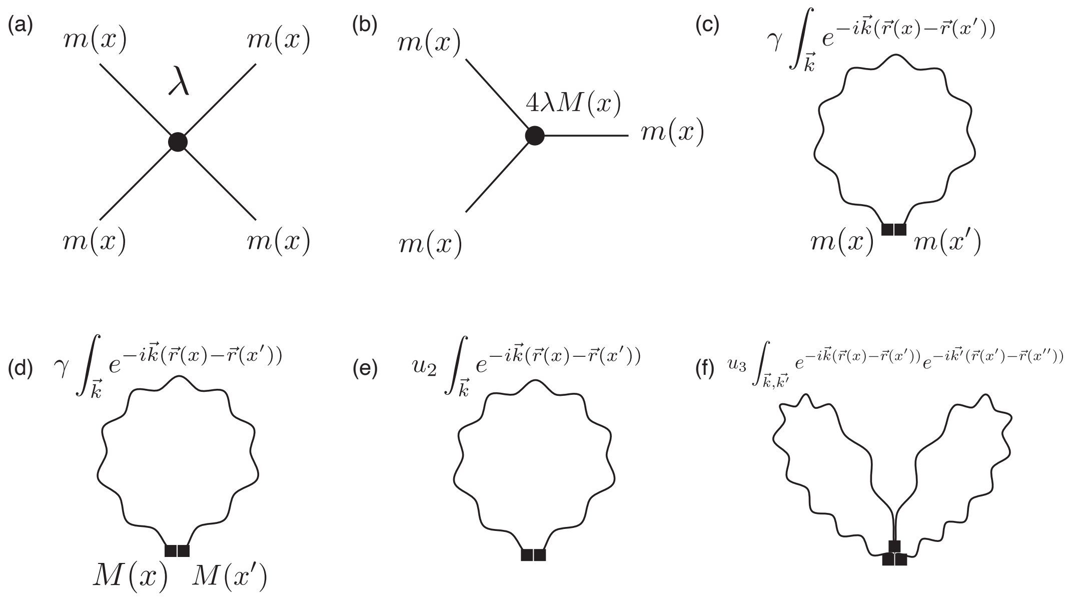


Figure S1. Interaction vertices generated by the vertices of the interacting part 2PI action. Solid lines correspond to the correlation function of the field $m(x)$. A single wavy line corresponds to the propagator of the two-body delta-function interaction $\delta(\vec{r}(x)-\vec{r}(x'))$, while a double wavy line corresponds to the propagator of the three-body delta-function interaction $\delta(\vec{r}(x)-\vec{r}(x'))\delta(\vec{r}(x')-\vec{r}(x''))$. The analytical expressions for the latter two interactions are shown in their three-dimensional Fourier representation. Vertices with mean-field insertions are indicated by the corresponding field symbol (e.g., $M(x)$). (a) Interaction vertex of the $m(x)^4$ term. (b) Interaction vertex generated by shifting the $m(x)^4$ term. (c) Interaction vertex of the polymer–field coupling. (d) Interaction vertex of the shifted polymer–field coupling. (e) Interaction vertex of the two-body polymer term. (f) Interaction vertex of the three-body polymer term.

The polymer-epigenetic field interaction produces two diagrams with expressions:

$$D_{\gamma 1} = \gamma \int_{x,x'} G_m(x,x')G_r(x,x')^{-3/2} \tag{S9}$$

$$D_{\gamma 2} = \gamma \int_{x,x'} M(x)M(x')G_r(x,x')^{-3/2}. \tag{S10}$$

The two-body interaction vertex graph corresponds to the expression:

$$D_{u_2} = u_2 \int_{x,x'} G_r(x,x')^{-3/2}. \tag{S11}$$

The three-body interaction vertex graph corresponds to the expression:

$$D_{u_3} = u_3 \int_{x,x',x''} [G_r(x,x')G_r(x',x'') - (G_r(x,x') - G_r(x,x'') + G_r(x',x''))^2]^{-3/2}. \tag{S12}$$

Combining these expressions, one obtains the lowest order form of $\Gamma_2\{M, \vec{R},\ G_m,\ G_r\}$.

The equations for $M^*$ and $G_m(x,x')$ are obtained by taking the derivative of the 2PI action with respect to $M(x)$ and $G_m(r,r')$. For the equations of $G_m(x,x')$, the corresponding diagrams are obtained from the ones entering $\Gamma_2\{M, \vec{R},\ G_m,\ G_r\}$ by truncating legs coming from the propagator $G_m(x,x')$ itself.

The dominant scaling equation for $G_r(x,x')$ is obtained by explicitly extracting the Fourier component of $G_r$, $\tilde{G}_r(q) = \int_x dx\ e^{iq\Delta x}G_r(\Delta x)$ in the expression for $D_{\gamma 1}$, $D_{u2}$ and $D_{u3}$ by $G_r(x,x') = \int_q dqe^{-iq(x-x')}\tilde{G}_r(q)$. We assume a scaling form for $\tilde{G}_r(q) = D|q|^{-2\alpha-1}$. Changing variables in the integrals over $x, x', x''$ in the expression for the diagrams in the 2PI as $qx = y, qx' = y', qx'' = y''$ and keeping only the terms proportional to powers of $|q|$ gives the dominant scaling equation Eq. (1).

##### B. Phase diagram calculation

The phase diagram in Fig. 1 is obtained by solving Eqs. (1),(2). To simplify the calculations, we are going to neglect the first-order contributions to the average $M^*$ and the correlation function $G_m$ in $\lambda$ and coming from the sixth-order term $m(x)^6$. We do not expect this approximation to change the picture qualitatively, as the mass correction at first order in $\lambda$ and the sextic interaction cause only a shift in the critical point position. We thus have to solve the set of coupled equations Eqs. (1), (2). We will carry out the integrals at a UV cut-off of 0.001 and an IR cut-off of 100. We will fix the values of $u_2,\ u_3,\ \lambda,\ \mu$ at $u_2 = 3.0,\ u_3 = 1.0, \lambda = 0.3,\ \mu = 0.5$ and vary the binding affinity $r_m$ and the polymer-field interaction strength $\gamma$. The transition line between the coil and globule polymer states is calculated by solving for those values of $\gamma$ for which $u_2 = -\gamma M^{*2}$ at fixed $r_m$ ($M^*$ fixed point of the flow at fixed parameters). The dashed horizontal line between the Globule phase/Disordered and Globule phase/Ordered is defined by the value of $\gamma$ for which the first derivative of Eq. (2) becomes negative: this line corresponds to the transition between a disordered and ordered state for the field $m$, in which, respectively, there are one and two stable minima.

##### C. Phase portraits in the $M - R$ plane

The phase portraits shown in Fig. 3 are obtained by considering the right-hand sides of Eqs. (1) and (2). To linear order, fluctuations around the fixed points are represented by the corresponding thermodynamic forces obtained by differentiating the 2PI action with respect to $M$ and $G_r$. We thus obtain the dynamical equations:

$$\Gamma_m \frac{dM}{dt} = -\partial_M \Gamma\{M, \vec{R}, G_m, G_r\} \tag{S13}$$

$$\Gamma_R \frac{dG_r(x,x')}{dt} = -\partial_{G_r} \Gamma\{M, \vec{R}, G_m, G_r\}, \tag{S14}$$

where $\Gamma_M$ and $\Gamma_R$ are the inverse mobilities of the $M$ and $G_r$ fields respectively. The time scales of the fields are defined as $T_M = \Gamma_m/\partial^2_M \Gamma$ and $T_R = \Gamma_R/\partial^2_R \Gamma$, where we have employed a quadratic expansion around the fixed points of Eqs. (S13),(S14).

For the $G_r$ dynamics, we evaluate the right-hand side of Eq. (1) at $q = 1/L$ with $L = 100$ and identify the intrachain distance $G_r(L) = R^2(L)$ from the Fourier representation $\widetilde{R^2}(q) = |q|^{-2\alpha-1}$. The $M$ component is obtained from the corresponding 2PI stationarity condition. To express this component as a function of $R(L)$, we invert the $M^* = 0$ relation between $R(L)$ and the second-virial coefficient in Eq. (1), obtaining $u_2 = u_2[R(L)]$. Substituting this relation into the stationary solution for $G_r$ yields the contact contribution $\int_{\Delta x} G_r(\Delta x)^{-3/2}$ to the mass of the $M$ field. For $R > 11.387$ (coil phase), the flow field of $M$ is set to that of the local theory, $-2\mu M^* - 4\lambda M^{*3} - 6M^5$. For visualization, each vector is normalized by its Euclidean norm; the arrows therefore indicate direction but not speed.

#### S3. MOLECULAR DYNAMICS SIMULATIONS

##### A. Model

We model the chromatin fiber as a bead-spring ring polymer of $N$ monomers, with arclength $L$. Each monomer $i$ is described by a three-dimensional position $\vec{r}_i \in \mathbb{R}^3$ and a continuous scalar field $m_i \in \mathbb{R}$ representing its local epigenetic state. The positions $\vec{r}_i$ and fields $m_i$ evolve according to overdamped Langevin equations,

$$\dot{\vec{r}}_i = \vec{F}_i^{\rm el} + \vec{F}_i^{\rm nb} + \boldsymbol{\xi}_i(t), \tag{S15}$$

$$\dot{m}_i = F_i^m + \eta_i(t), \tag{S16}$$

where $\boldsymbol{\xi}_i$ and $\eta_i$ are independent Gaussian white-noise terms with zero mean and identical variance,

$$\langle \xi_i^\alpha(t)\, \xi_j^\beta(t')\rangle = 2D\, \delta_{ij}\, \delta^{\alpha\beta}\, \delta(t-t'), \qquad \langle \eta_i(t)\, \eta_j(t')\rangle = 2D\, \delta_{ij}\, \delta(t-t'), \tag{S17}$$

with diffusion constant $D$ controlling the amplitude of thermal fluctuations for both positional and epigenetic degrees of freedom. The epigenetic diffusion constant along the chain contour is set to $D_m = 0$, so that spreading of the epigenetic field occurs exclusively through the three-dimensional polymer contacts encoded in $F_i^m$. In simulations, we used rescaled versions of the parameters introduced in the theory.

##### B. Bonded interactions

Adjacent monomers along the chain are connected by Hookean springs. The elastic force on monomer $i$ is

$$\vec{F}_i^{\text{el}} = -k_{\text{el}} \sum_{j\in\{i-1,\, i+1\}} \frac{|\vec{r}_{ij}| - r_0}{|\vec{r}_{ij}|}\,\vec{r}_{ij}, \tag{S18}$$

where $\vec{r}_{ij} = \vec{r}_i - \vec{r}_j$, $r_0$ is the equilibrium bond length, and $k_{\text{el}}$ is the spring constant. All inter-monomer distances are evaluated using minimum-image periodic boundary conditions to handle chains that span the simulation box of side $L_{\text{box}} = 100$.

##### C. Non-bonded excluded-volume repulsion

Short-range steric repulsion between nonbonded monomers (i.e., monomers that are not nearest neighbors along the chain) is modeled by a modified Weeks–Chandler–Andersen (WCA) potential, which retains only the repulsive branch of the Lennard-Jones interaction. The corresponding force reads

$$\vec{F}_{ij}^{\text{WCA}} = 48\varepsilon \left[ \left(\frac{\sigma^2}{r_{ij}^2}\right)^5 - \frac{1}{2}\left(\frac{\sigma^2}{r_{ij}^2}\right)^2 \right] \vec{r}_{ij} ds, \qquad r_{ij} < 2^{1/6}\sigma, \tag{S19}$$

and vanishes for $r_{ij} \geq 2^{1/6}\sigma$. The corresponding potential is continuous at the cutoff and diverges as $r_{ij} \to 0$. Here, $ds = L/N$ is the monomer arc-length spacing. This renders the chain self-avoiding. During the initial equilibration phase (see below), the WCA potential is replaced by a softer cosine repulsion,

$$F^{\,\text{soft}}(r_{ij}) = A\cos\!\left(\frac{\pi}{2}\frac{r_{ij}}{r_c}\right), \qquad r_{ij} < r_c = 2^{1/6}\sigma, \tag{S20}$$

which remains finite at $r_{ij} = 0$ and avoids numerical blow-up when monomers overlap during the disentanglement of the initial configuration.

##### D. Polymer–epigenetic field coupling

Non-bonded monomers interact through a field-mediated Yukawa-type potential. The spatial force on monomer $i$ due to monomer $j$ is

$$\vec{F}_{ij}^{\text{att}} = \frac{\gamma\,\Lambda\, e^{-\Lambda r_{ij}}}{r_{ij}}\, m_i\, m_j\, \vec{r}_{ij} ds, \tag{S21}$$

where $\gamma$ is the coupling strength and $\Lambda^{-1}$ is the interaction range. This force is attractive when $m_i$ and $m_j$ share the same sign, promoting spatial clustering of monomers with identical epigenetic identity, and repulsive otherwise. The total nonbonded spatial force on monomer $i$ is

$$\vec{F}_i^{\text{nb}} = \sum_{j\neq i} \left( \vec{F}_{ij}^{\text{WCA}} + \vec{F}_{ij}^{\text{att}} \right). \tag{S22}$$

The epigenetic field of monomer $i$ evolves under the combined effects of an external writing field, local relaxation and nonlinear saturation, and Yukawa coupling to neighboring fields:

$$F_i^m = r_m - \mu\, m_i - \lambda\, m_i^3 - m_i^5 - \gamma \sum_{j\neq i} e^{-\Lambda\, r_{ij}}\, m_j. \tag{S23}$$

Here, $r_m$ is the external writing field, $\mu$ controls linear relaxation, and the cubic and quintic terms limit the field amplitude; together with contact-mediated feedback, these terms can generate two stable epigenetic states. The last sum implements a positive-feedback reader–writer mechanism: spatially proximate monomers with a nonzero field bias the field of monomer $i$ toward their own sign, modeling the enzymatic spreading of histone modifications. Since the one-dimensional diffusion constant for the field is set to $D_m = 0$, this three-dimensional contact-mediated coupling is the sole mechanism by which epigenetic information propagates along the chain.

##### E. Adaptive time-stepping and spatial neighbor search

To maintain numerical stability under the large force gradients generated by the WCA potential, the simulation employs an adaptive time step $\delta t$ computed at each iteration as

$$\delta t = \min\left( dt_{\max},\ \frac{\delta x_{\text{thresh}}}{\max_i |\vec{F}_i^{\text{el}} + \vec{F}_i^{\text{nb}}|} \right), \tag{S24}$$

where $dt_{\max}$ is the maximum allowed time step and $\delta x_{\text{thresh}}$ is the maximum allowed monomer displacement per step. The noise amplitude is rescaled accordingly as $\sqrt{2D\,\delta t}$ at each step, consistently for both positional and epigenetic noise.

Non-bonded force evaluation is accelerated by a cell-list (linked-cell) algorithm: the simulation box is divided into cubic cells of side equal to the cutoff radius (set to 5.0), and the neighbor search is restricted to the $3 \times 3 \times 3 = 27$ cells surrounding each monomer. The cell list is rebuilt every $n_{\text{list}} = 5$ time steps. All kernels are implemented in CUDA using the Thrust library and executed on GPU, with monomer data stored as `float4` vectors $(x, y, z, m)$ to exploit coalesced memory access. The polymer and field configurations were saved every $dt_{\text{save}} = 0.01$.

##### F. Initialization protocol

The simulation proceeds through three sequential phases designed to produce a well-equilibrated initial configuration before the full coupled dynamics are activated.

*Phase 1 – Gaussian chain relaxation.* The chain is initialized as a random walk and evolved as a phantom (non-self-avoiding) Gaussian chain for time $T_1 = 10.0$, with only elastic bonds active and no non-bonded interactions. This allows the chain to relax any residual stretching from the initial configuration while preserving chain connectivity.

*Phase 2 – Self-avoiding walk equilibration.* The soft cosine repulsion is activated and the chain is evolved for an additional time $T_2 = 40.0$. Monomer overlaps inherited from the random-walk initialization are eliminated without the risk of numerical instabilities, and the chain adopts a proper self-avoiding conformation.

*Phase 3 – Full coupled dynamics.* The WCA repulsion replaces the soft repulsion, the attractive Yukawa coupling is activated, and the epigenetic-field dynamics [Eq. (S23)] are switched on with the chosen values of $r_m$, $\mu$, $\lambda$, $\gamma$, and $\Lambda$. If the constant-field initialization option is selected, the epigenetic fields are first set to a prescribed uniform average value $m_i = m_0$ before the dynamics are activated, providing a controlled starting condition. The system is then evolved until a steady state is reached at time $T$.

##### G. Reprogramming protocol

We model reprogramming by adding the external field derived from the reprogramming free-energy term introduced in Sec. S1 to the equation of motion for $m$. This term is given by $h_{\text{ext}} = -h_0 \operatorname{sgn}(m_i)$. The field is applied at simulation time $t = 100$, when we set $h_0 = 7.0$ from $h_0 = 0$. This occurs after the three initialization phases described in Sec. S3 F, so that the polymer has reached its equilibrium distribution. The three regimes probed in Fig. 3 correspond to induction durations $t_{\text{ind}} = 0.2,\ 4,\ 20$. For the figure shown in the main text, Fig. 3, we simulated a chain of $N = 1024$ monomers. The parameters chosen are shown in Table I.

##### H. Analysis of simulation trajectories

To further quantify the memory and plasticity properties of the epigenetic field along the separatrix, we run simulations at different protocol induction durations for a polymer of size $N = 512$ and for a protocol of strength $h_0 = 3$, all other parameters kept fixed as shown in Table I.

To quantify memory of the initial epigenetic pattern independently of changes in its spatial mean, we calculated the absolute connected epigenetic overlap. Writing $\delta m_i(t) = m_i(t) - M(t)$, where $M(t) = N^{-1} \sum_i m_i(t)$, we defined

$$|\rho_m(t, t_1)| = \left| \frac{\sum_i \delta m_i(t_1) \delta m_i(t)}{\sqrt{\sum_i \delta m_i^2(t_1) \sum_i \delta m_i^2(t)}} \right| .$$

| Parameter list for reprogramming simulation | |
|---|---|
| Parameter | Value |
| $\gamma$ | -5.0 |
| $r_m$ | 0.0 |
| $\mu$ | 1.0 |
| $\Lambda$ | 1.0 |
| $\lambda$ | 0.1 |
| $N$ | 1024 |
| $L$ | 102.4 |
| $r_0$ | 1.122 |
| $\sigma$ | 1.0 |
| $\varepsilon$ | 1.0 |
| A | 30.0 |
| $k_{el}$ | 500 |
| $h_0$ | 7.0 |
| $D$ | 1 |
| $\delta x_{\text{thresh}}$ | 0.01 |
| $dt_{\max}$ | 0.0001 |

Table I. Table of parameters used for the simulation shown in Fig. 3.

Here, $t_1$ denotes the initial pre-induction configuration. Subtraction of the instantaneous mean ensures that $\rho_m$ measures memory stored in spatial fluctuations rather than persistence of the uniform order parameter $M$. Taking the absolute value identifies recovery of the initial fluctuation pattern modulo a global inversion of the epigenetic marks.

To determine whether the polymer retains its locus-specific three-dimensional organization, rather than merely remaining compact, we calculated a conformational contact overlap. For nonlocal monomer pairs with ring contour separation $s_{ij} = \min(|i-j|, N-|i-j|) > 2$, we defined the weighted contact kernel

$$K_{ij}(t) = e^{-\Lambda r_{ij}(t)}\Theta(r_c - r_{ij}(t)),$$

using minimum-image distances, $\Lambda = 1$, and $r_c = 5$. The generic dependence on contour separation was removed through $\delta K_{ij}(t) = K_{ij}(t) - \langle K_{kl}(t)\rangle_{s_{kl}=s_{ij}}$. The normalized conformational overlap was then

$$Q_{\text{conf}}(t,t_1) = \frac{\sum_{i<j} \delta K_{ij}(t_1)\delta K_{ij}(t)}{\sqrt{\sum_{i<j} \delta K_{ij}^2(t_1) \sum_{i<j} \delta K_{ij}^2(t)}}.$$

Thus $Q_{\text{conf}}(t_1,t_1) = 1$, while loss of the specific initial contact network drives the overlap toward its label-shuffled baseline even if the radius of gyration remains small.

To quantify the instantaneous plasticity of the mean epigenetic state while avoiding the divergence of a static inverse-Hessian response near the separatrix, we calculated a finite-time susceptibility with response horizon $\Delta = 0.5$. At each saved configuration, the polymer conformation was held fixed, and the epigenetic Hessian $H(t)$ was diagonalized, $H\mathbf{v}_\alpha = \lambda_\alpha \mathbf{v}_\alpha$. The response to a weak spatially uniform field was then defined as

$$\chi_M(t;\Delta = 0.5) = \sum_\alpha w_\alpha(t) \frac{1 - e^{-0.5\lambda_\alpha(t)}}{\lambda_\alpha(t)}, \qquad w_\alpha(t) = \left| \mathbf{v}_\alpha(t) \cdot \frac{\mathbf{1}}{\sqrt{N}} \right|^2,$$

with the continuous limit $(1 - e^{-0.5\lambda})/\lambda \to 0.5$ for $\lambda \to 0$. The weights $w_\alpha$ select modes that couple to the spatially averaged order parameter $M$, so a soft eigenmode contributes strongly only when it has a substantial uniform component. Thus, $\chi_M(t;0.5)$ measures the predicted change of $M$ generated by a weak uniform perturbation acting for half a simulation-time unit, providing a finite and dynamically meaningful measure of epigenetic susceptibility at each point along the trajectory.

##### I. Contact probability and connected correlation functions from molecular dynamics simulations

To verify the predicted relationship between the connected field correlation function and the polymer contact probability, we calculated both quantities from molecular dynamics simulations. We considered three interaction strengths, $\gamma_{\mathrm{SAW}} = -4.0$, $\gamma_1 = -5.0$, and $\gamma_2 = -7.0$. All other parameters were set as in the perturbation protocol (Table I). The first value corresponds to a self-avoiding-walk phase, whereas the latter two correspond to collapsed globules. For $\gamma_{\mathrm{SAW}}$, we simulated the system for a total time $T = 1500$, including phases 1 and 2 of the initial equilibration procedure. For $\gamma_1$ and $\gamma_2$, we simulated the system until $T = 1000$. At each time $t$, we calculated $G_m(x, x', t)$ from $\widetilde{m}(q,t)\widetilde{m}(-q,t)$ after setting the zero mode $\widetilde{m}(0,t)$ to zero, thereby removing the spatial mean. Polymer contacts were counted within a radius of 1.0. The mean connected correlation function and the contact probability $P(s)$ were obtained by averaging samples collected after $T = 250$. Results are shown in Fig. S2. For $\gamma = -4.0$, the decay of the contact probability is consistent with self-avoiding-walk statistics, $P(s) \propto s^{-2.1}$. The connected field correlation function decays with the same scaling. At $\gamma = -5.0$, we instead observe fractal-globule scaling, $P(s) \propto |x - x'|^{-1}$, with the connected correlation function scaling accordingly. At the higher interaction strength $\gamma = -7.0$, the connected correlation function decays more slowly because of the longer correlation length.

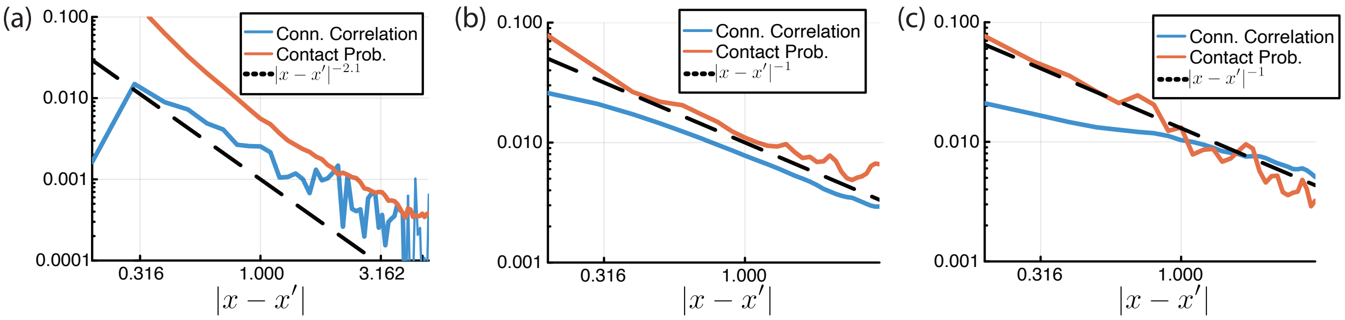


Figure S2. Connected correlation functions and contact probability from molecular dynamics simulations at three different values of $\gamma$. The dashed line indicates the expected scaling for the corresponding polymer phase. (a) $\gamma = -4.0$. (b) $\gamma = -5.0$, (c) $\gamma = -7.0$.

#### S4. CALCULATION OF CORRELATION FUNCTIONS IN ESC-2I AND MEF CELLS

The datasets used for the data analysis in the main text are shown in Table II.

For the MEF data, we back-transformed the reported $\log_2$ H3K9me3 read counts and averaged them in 200-bp bins to reduce read noise. Because the multiscale organization of chromatin can dominate whole-genome correlation functions, we restricted the analysis to H3K9me3 signals within topologically associating domains (TADs) [50]. To reduce heterogeneity among genomic contexts, we further averaged correlation functions only across TADs with similar epigenetic signatures. Compartments and TADs were identified from Hi-C data using the HiCExperiment and HiContacts R packages. For MEFs, compartments and TADs were calculated at resolutions of 200 kb and 40 kb, respectively; for 2i ESCs, the corresponding resolutions were 200 kb and 50 kb. For each TAD and compartment, we calculated the mean signals of the selected epigenetic marks (Table II), focusing on repressive marks characteristic of compact chromatin. We then grouped TADs by these mean signals using the R $k$-means implementation. This procedure yielded four clusters with distinct epigenetic signatures for each cell type (Figs. S3 and S4). Finally, we calculated connected H3K9me3 correlation functions in 1-Mb windows within TADs belonging to the same cluster.

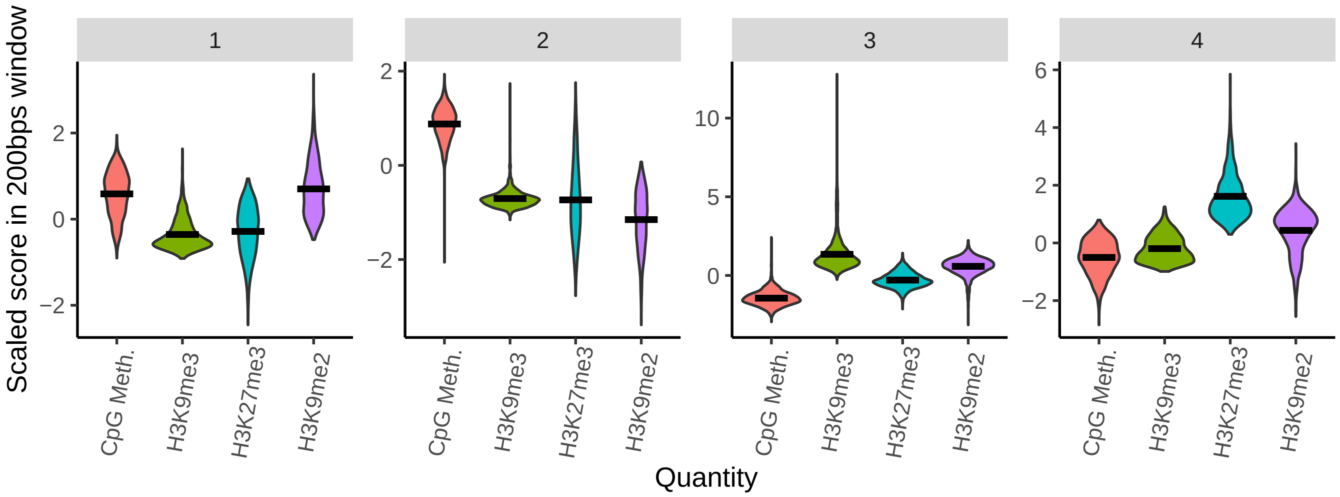


Figure S3. Distribution of epigenetic marks within the detected MEF TAD clusters. The $y$ axis represents the scaled mean signal within each TAD, and the $x$ axis indicates the corresponding mark.

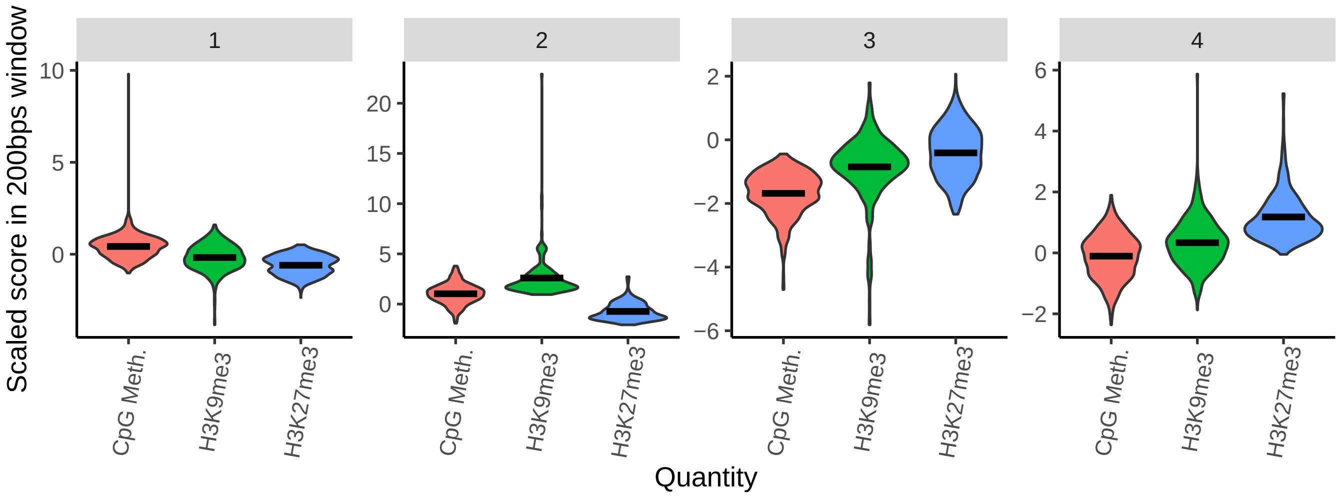


Figure S4. Distribution of epigenetic marks within the detected 2i-ESC TAD clusters. The $y$ axis represents the scaled mean signal within each TAD, and the $x$ axis indicates the corresponding mark.

| GEO accession numbers for data used | |
|---|---|
| Data | GEO accession number |
| MEF H3K9me3 | GSM6001986 |
| MEF H3K9me3 | GSE200011 |
| MEF H3K27me3 | GSM6001989 |
| MEF DNA methylation | GSM6001978 |
| MEF Hi-C | GSE200012 |
| 2i Hi-C | GSE119171 |
| 2i DNA methylation | GSM1027572 |
| 2i H3K27me3 | GSM590116 |
| 2i H3K9me3 | GSM850407 |

Table II. Table of GEO accession numbers for the data used in the calculations of the correlation functions.
